# Endothelial β-arrestins Regulate Mechanotransduction by the Type II Bone Morphogenetic Protein Receptor in Primary Cilia

**DOI:** 10.1101/2022.02.04.479175

**Authors:** Saejeong Park, Zhiyuan Ma, Georgia Zarkada, Irinna Papangeli, Sarin Paluri, Nour Nazo, Xinyu Xiong, Felix Rivera-Molina, Derek Toomre, Sudarshan Rajagopal, Hyung J. Chun

## Abstract

**Rationale:** Modulation of endothelial cell behavior and phenotype by hemodynamic forces involves many signaling components, including cell surface receptors, intracellular signaling intermediaries, transcription factors, and epigenetic elements. Many of the signaling mechanisms that underlie mechanotransduction by endothelial cells are inadequately defined.

**Objective:** We sought to better understand how β-arrestins, intracellular proteins that regulate agonist-mediated desensitization and integration of signaling by transmembrane receptors, may be involved in the endothelial cell response to shear stress.

**Methods and Results:** *In vitro* studies with primary endothelial cells subjected to β-arrestin knockdown, and *in vivo* studies using mice with endothelial specific deletion of β-arrestin 1 and β-arrestin 2 were conducted. We found that β-arrestins are localized to primary cilia in endothelial cells, which are present in subpopulations of endothelial cells in relatively low shear states. Recruitment of β-arrestins to cilia involved its interaction with IFT81, a component of the flagellar transport protein complex in the cilia. β-arrestin knockdown led to marked reduction in shear stress response, including induction of *NOS3* expression. Within the cilia, β-arrestins were found to associate with the type II bone morphogenetic protein receptor (BMPR-II), whose disruption similarly led to an impaired endothelial shear response. β-arrestins also regulated Smad transcription factor phosphorylation by BMPR-II. Mice with endothelial specific deletion of β-arrestin 1 and β-arrestin 2 were found to have impaired retinal angiogenesis.

**Conclusion:** We have identified a novel role for endothelial β-arrestins as key transducers of ciliary mechanotransduction that play a central role in shear signaling by BMPR-II and contribute to vascular development.

**NOVELTY AND SIGNIFICANCE:** 

**What Is Known?:**

- Endothelial cells respond to flow-induced shear stress with biochemical changes, such as phosphorylation of endothelial nitric oxide synthase, that promote morphological changes, such as cell alignment.
- The endothelial response to shear stress can involve primary cilia, microtubule-based sensory organelles that detect extracellular stimuli and generates intracellular signals.
- The specific ciliary signaling pathways that regulate endothelial mechanotransduction have not been fully elucidated.

**What New Information Does This Article Contribute?:**

- β-arrestins directly interact with the ciliary protein intraflagellar transport protein 81 (IFT81), which is present in the primary cilia of endothelial cells, and are required for the morphological response to flow-induced shear stress.
- β-arrestins regulates type II bone morphogenetic protein receptor signaling, which is required for the endothelial response to shear stress, and is required for the phosphorylation of Smad transcription factors.
- β-arrestins are required for endothelial nitric oxide synthase-mediated flow-induced shear stress response in endothelial cells.
- Endothelial cell-specific knockout of β-arrestins results in abnormal vascular development, with a loss of vessel length and branchpoints.

## INTRODUCTION

Sensing of the environment by endothelial cells (ECs) is a complex phenomenon that regulates the development, physiology and pathophysiology of vascular and lymphatic structures. These environmental stimuli include vasoactive mediators and mechanical stress that are central to EC biology^1^. ECs are under constant mechanical stress and sense variations in blood flow-induced shear stress^2^. Shear stress induces the expression of thousands of genes in ECs, with differing effects on cellular phenotype depending on the direction and the magnitude of the flow pattern. For example, unidirectional laminar flow promotes elongation and alignment of ECs^3-5^ and the expression of anti-inflammatory genes^6^, while turbulent flow stimulates disorganized cell proliferation with the expression of inflammatory and oxidative genes^7^. The underlying mechanism for such differential response remains to be fully elucidated. These phenotypic changes in ECs are controlled by their sensing and transduction of mechanical signals during vascular remodeling. Multiple signaling pathways are regulated by shear stress, include MAP kinases, Rho family GTPases, reactive oxygen species, nitric oxide (NO), and receptors such as G protein-coupled receptors (GPCRs), bone morphogenetic protein receptors (BMPRs), and vascular endothelial growth factor receptors^8^.

Primary cilia are microtubule-based organelles found in most mammalian cell types that sense extracellular stimuli^1, 9^, using sensors of the external environment such as GPCRs, receptor tyrosine kinases, and BMPRs^10^. The primary cilia contain proteins critical for protein trafficking, structure and signaling, such as adenylyl cyclases, protein phosphatases and protein kinases^11^. Primary cilia have been shown to respond to blood flow and are required for flow sensing^12, 13^. Receptors have been shown to be specifically trafficked to the primary cilia^14^, resulting in a unique signaling environment with increased levels of local second messengers such as cyclic AMP^15^ that result in distinct outcomes from extraciliary messengers^16^. Flow or shear stress can activate the primary cilium, triggering calcium signaling and nitric oxide production^9^. Primary cilia play a role in vascular development, with expression in mouse retinal endothelia where they promote vessel stabilization through increased sensitivity of the BMP-Alk1 pathway^17^. They have been shown to be essential for appropriate vascular patterning and responsible for enhancing BMPR-II signaling^17^. Diseases of the primary cilia, known as ciliopathies, have important cardiovascular phenotypes related to development, hypertension, and atherosclerosis^18^. However, the precise signaling pathway of mechanotransduction by primary cilia have not been fully elucidated, despite their physiological importance in vascular biology.

With their known role in trafficking ciliary proteins and regulating GPCR signaling^19^, we sought to elucidate the role of β-arrestin adaptor proteins in ECs. We conducted an unbiased interactome screen to identify proteins physically associated with β-arrestins, and found multiple ciliary proteins to be associated with β-arrestin. We found disruption of β-arrestin expression in ECs led to marked impairment of EC response to shear stress. Moreover, we found that ciliary shear response is at least in part mediated by BMPR-II, whose downstream signaling cascade also requires β-arrestin. Lastly, mice with conditional, endothelial specific deletion of β-arrestin 1 and β-arrestin 2 were found to have marked impairment in their retinal angiogenesis, providing in vivo evidence for the role of β-arrestin in EC function and vascular development.

## METHODS

### Materials

Human recombinant BMP-2 and BMP-4 were purchase from R&D Systems (Minneapolis, MN). Anti-Myc and anti-HA antibodies were purchased from Cell Signaling Technology (Danvers, MA). FLAG-β-arrestin 1 and 2 plasmids and rabbit polyclonal anti-β-arrestin1 and 2 antibodies (A1CT and A2CT) were provided by Dr. Robert Lefkowitz (Duke University, Durham, NC). Plasmids encoding HA or Myc-tagged BMPR2, ALK3 and ALK6 were provided by Dr. Gerard Blobe (Duke University, Durham, NC), the HA and Myc-tagged S532X BMPR2 plasmids were provided by Dr. Nicholas Morrell (Cambridge University, Cambridge, UK), the bre-luc reporter was provided by Dr. Peter ten Dijke (Netherlands Cancer Institute, Amsterdam, the Netherlands) and parental and β-arrestin1/2 CRISPR KO HEK293 cells were provided by Asuka Inoue (Tohoku University, Sendai, Japan).

### Cell culture

Human umbilical vein endothelial cells (HUVECs) (Lonza) were cultured in EGM-2 medium (Lonza) at 37 °C in a 5% CO_2_ incubator. The HUVECs (passages 4–9) were grown to 80-90 % confluence for experimental treatments. HEK293 cells (WT and β-arrestin1/2 KO cells) were grown in MEM with 10% FBS and 1% penicillin/streptomycin. Starvation medium consisted of MEM supplemented with 0.1% BSA, 10 mM HEPES and 1% penicillin/streptomycin.

### Transient Transfection and siRNA knockdown

Transient transfection of plasmids was performed with Fugene HD (Promega) according to the manufacturer’s instructions. For RNA silencing in HUVECs, short interfering RNAs (siRNAs) for both targeting sequences and non-targeting controls (GE Dharmacon, 50 nM, 72 h) were transfected with Lipofectamine RNAiMAX (Invitrogen) in HUVECs according to the manufacturer’s instructions. siRNA knockdown of β-arrestins in HEK293s was performed as previously described^20^. siRNA sequences targeting β-arrestin1 (5’0AAAGCCUUCUGCGCGGAGAAU-3’) and β-arrestin2 (5’-AAGGACCGCAAAGUGUUUGUG-3’) along with a control non-silencing RNA duplex (5’0AAUUCUCCGAACGUGUCACGU-3’) were used. Cells were transfected using the GeneSilencer transfection reagent (Genlantis) as previously described^20^.

### Shear stress condition and cell alignment assay

To subject HUVECs to orbital shear stress, calculated at 10.7 dynes/cm^2^, an orbital shaker positioned inside the cell incubator was used. All shear stress experiments were performed for 24 h. To observe cell alignment under shear stress, the HUVECs were grown on 6-mm culture dishes and the cells 2-mm away from the edge of the well were taken as images using a brightfield microscope.

### Proteomic analysis

HUVECs were subjected to 10.7 dynes/cm^2^ shear stress for 24 hours (shear samples) or maintained under static conditions (static samples), washed once in PBS and collected with NP-40 lysis buffer (Boston Bioproducts) with protease and phosphatase inhibitors (Roche). The cell lysates from the shear and static samples were immunoprecipitated separately with rabbit anti-β-arrestin1/2 antibody (CST) overnight at 4 °C and protein A/G agarose beads (Santa-Cruz) for 2 h with rotation at 4 °C. After two rinses in lysis buffer and one in PBS, the shear and static samples were profiled by tandem mass spectrometry and ultraperformance liquid chromatography (Bioproximity, LLC). Results were reported for proteins observed in the shear but not the static conditions.

### Immunofluorescence staining

HUVECs grown on 8-well cell culture slides (Mat-Tek) coated with Fibronectin (Corning, Sigma-Aldrich) were washed with warmed PBS (37 °C) twice, fixed in warm 4% paraformaldehyde (Thomas Scientific, 37 °C) for 5 min and washed with PBS 3 times. The cells were permeabilized with 1% Triton X-100 in PBS for 5 min and washed again with PBS 3 times. The cells were blocked with 1% bovine serum albumin (Sigma-Aldrich) in PBS for 30 min. Anti-acetylated-tubulin (T6793, Sigma-Aldrich, 1:1000), anti-Arl13B (17711-1-AP, Proteintech, 1:500), anti-IFT81 (11744-1-AP, Proteintech, 1:500) and anti-BMPR-II (612292, BD Biosciences, 1:500) antibodies were incubated overnight at 4°C, and then detected with Alexa Fluor 568 donkey anti-mouse IgG (A10037, Invitrogen, 1:500), Alexa Fluor 568 goat anti-rabbit IgG (A11011, Invitrogen, 1:200), Alexa Fluor 488 goat anti-mouse IgG (A11001, Invitrogen, 1:200) and/or Alexa Fluor 647 goat anti-rabbit IgG (A21244, Invitrogen, 1:200). 0.5 µg/ml DAPI (Sigma-Aldrich) was used to stain the nuclei. The cells were mounted using Vectashield (Vector Labs) and imaged using fluorescence microscopy.

### Confocal microscopy of β-arrestin 2 recruitment

HEK293 cells were transiently transfected with vectors encoding red fluorescent protein (RFP)-labeled β-arrestin2, BMPR2 and either of its co-receptors ALK3 or ALK6. 2 µg of each plasmid was combined with Fugene transfection reagent (Roche Applied Science, Mannheim, Germany) at a ratio of 5 µl of transfection reagent to 1 µg DNA in 500 µl of MEM and incubated at room temperature for 30 minutes prior to addition to a 10 cm dish of HEK293 cells at 50% confluence. 24 hours after transfection, the cells were split into 35 mm glass-bottom dishes (MatTek, Ashland, MA). After a further 24-48 hours of growth, the cells were starved for at least 1 hour prior to fixation or stimulation with 50 ng/ml BMP-2 for 30 minutes followed by fixation. Cells were fixed in 1 ml 4% paraformaldehyde in PBS for 10-15 min. RFP-labeled β-arrestin2 was visualized on a Marianas spinning disk confocal microscope (Intelligent Imaging Innovations, Inc., Denver, CO).

### Western blot in HUVECs

β-arrestin1 and β-arrestin2 siRNA (50 nM) or non-targeting siRNA (50 nM) were transfected into HUVECs for 24 h and the proteasome inhibitor MG132 (0.2 µM, Fisher Scientific) was added to the HUVECs for 48 h. The HUVECs were washed with PBS and lysed in NP-40 lysis buffer (Boston Bioproducts) including protease and phosphatase inhibitor (Roche). The cell lysates were incubated rotated with rabbit anti-eNOS antibody (CST, 1:50) at 4 °C for overnight. Protein A/G agarose beads (Santa-Cruz) were added to immunoprecipitated samples and rotated for 2 h with rotation at 4°C. After twice washes in lysis buffer and once in PBS at 4 °C, the agarose beads were boiled in Laemmli SDS sample buffer (Alfa Aesar) for western blot analysis with rabbit anti-eNOS antibody (CST, 1:1000), Ubiquitin antibody (Santa Cruz, 1:200), β-arrestin antibody (CST, 1:1000) and GAPDH antibody (CST, 1:1000).

### Protein extraction and western blotting

HUVECs were lysed in NP40 lysis buffer (Boston Bioproducts) including cOmplete™ Mini EDTA-free Protease Inhibitor Cocktail (Roche; Sigma-Aldrich) and PhosSTOP™ Phosphatase Inhibitor tablets (Roche; Sigma-Aldrich). The lysed supernatant was obtained by centrifugation at 4°C for 10 min at 13,000 rpm and the total protein content was quantified by a Micro BCA Protein assay kit (Thermo Scientific). The proteins were boiled in Laemmli SDS sample buffer (Alfa Aesar) for 10 min and separated by Mini-Protean TGX Gels 4-20 % (Bio-Rad), transferred onto an Immobilon-PSQ PVDF Membrane (EMD Millipore). Membranes were probed with antibodies against β-arrestin1/2, phospho-eNOS (Ser1177), total-eNOS, Myc-tag, Erk5, GAPDH (CST, 1:1000), IFT81, IFT74 (Proteintech, 1:1000), GFP (Sigma-Aldrich) and BMPR-II (BD Bioscience, 1:1000). Bound primary antibodies were detected by secondary anti-rabbit or anti-mouse IgG-HRP (CST, 1:2000). Blots were developed with Supersignal West Femto Maximum Sensitivity Substrate (Thermo Scientific).

### Co-immunoprecipitation (CoIP) of β-arrestin1 and IFT81

After co-transfection with GFP-tagged β-arrestin1 or GFP control constructs and Myc-tagged IFT81 constructs, HEK293 cells were washed with PBS and lysed in NP-40 lysis buffer (Boston Bioproducts) including protease and phosphatase inhibitor (Roche). The cell lysates were incubated/rotated with mouse anti-GFP antibody (Sigma-Aldrich, 1:20) at 4 °C overnight. Protein A/G agarose beads (Santa-Cruz) were added to immunoprecipitated samples and rotated for 2 h with rotation at 4 °C. After two washes in lysis buffer and one in PBS at 4 °C, the agarose beads were boiled in Laemmli SDS sample buffer (Alfa Aesar) for western blot analysis with rabbit anti-Myc antibody (CST, 1:1000), mouse anti-GFP antibody (Sigma-Aldrich, 1:1000) and GAPDH antibody (CST, 1:1000).

### CoIP of BMPR-II and β-arrestins

For CoIP in HEK293s, cells were grown in 6-well plates to 50% confluency and co-transfected with Myc-BMPR2 or Myc-BMPR2-S532X and HA-β-arrestin1/2 using Lipofectamine 2000 (Invitrogen). After 48 h of transfection, cells were scraped and lysed in lysis buffer of Pierce Crosslink IP Kit (Thermo/Pierce) with complete protease inhibitor cocktail tablet (Roche Applied Sciences). Cell lysates were incubated overnight at 4°C with anti-Myc-sepharose made with anti-Myc antibody (Cell signaling) and Pierce Crosslink IP Kit (Thermo/Pierce). Sepharose beads were then eluted with 0.5 mg/ml c-Myc peptide (Genscript) in TBS. The elution samples were then detected by immunoblotting.

### Bre-luc reporter

The bre-luc luciferase reporter of Smad activity was obtained as a gift from Peter ten Dijke^21^. A 10 cm dish of HEK293 cells of ∼ 50% confluence was transiently transfected with 2 µg of the reporter and 20 µg of control, β-arrestin1 or 2 siRNA with 60 µl of Lipofectamine 2000 (see above). After 24 hours, cells were split into 96 well plates at 50,000-100,000 cells/well. 24 hours later, cells were starved for 4 hours and treated overnight with 50 ng/ml BMP-2. Cells were then washed with room temperature PBS. Next, 80 µL of a coelenterazine-h/HBSS solution (3 µM coelenterazine-h) was added. Luminescence was then assayed with a NOVAstar plate reader (BMG Labtech, Cary, NC).

### Receptor Biotinylation

HEK293 cells transfected with expression plasmids for HA-BMPR2 and FLAG-β-arrestin1/2 were surface biotinylated at 4 °C for 20 minutes using EZ-link Sulfo-NHS-SS-Biotin (Pierce/Thermo). Unreacted biotin was quenched with 50 mM glycine in PBS at 4 °C. For the endocytosis assay, cells were incubated with 100 ng/ml BMP-2 at 37 °C for indicated time and remaining surface biotin was cleaved on ice with glutathione cleavage buffer (50 mM glutathione, 75 mM NaCl, 75 mM NaOH, 10 mM EDTA, 1% BSA in H_2_O). Excess glutathione was then quenched at 4 °C in iodoacetamide buffer (5 mg/ml iodoacetamide in PBS). Cells were then scraped and lysed in glycerol lysis buffer and biotinylated receptors were isolated by NeutrAvidin agarose (Pierce/Thermo) precipitation and detected by immunoblotting.

### Animals

The *Arrb1*^*fl/fl*^, *Arrb2* ^*fl/fl*^, and *Cdh5(PAC)-CreERT2* mice have been previously described ^22-24^. To generate conditional, endothelial-specific *Arrb1/Arrb2*-deficient mice (*Arrb1/Arrb2* ECKO), *Cdh5(PAC)-CreERT2* mice were crossbred with *Arrb1*^*fl/fl*^ and *Arrb2* ^*fl/fl*^ mice. To induce recombination of the *Arrb1/Arrb2* alleles with the *Cdh5-CreERT2* driver, mice were injected intraperitoneally at P1, P2 and P3 with 50 μg of Tamoxifen as previously described ^25^. All mice used in the present study were backcrossed to the *C57BL/6* strain background for at least 5 generations. All animal experimental protocols were approved by the Institutional Animal Care and Use Committee at Duke University Medical Center.

### Retinal angiogenesis analyses

Retinas dissected from p7 mice were washed by cold PBS once, pre-fixed in 4 % paraformaldehyde for 15 min in room temperature, washed by cold PBS once, fixed in cold methanol and stored at -20 °C for overnight. The retinas were washed in PBS for 10 min 3 times, re-fixed in 4 % paraformaldehyde for 10 min at room temperature and washed in PBS for 10 min 3 times. The retinas were permeabilized and blocked with 1% BSA and 0.5% Triton X-100 in PBS for 1 h at room temperature, followed by incubation with anti-eNOS primary antibody (32027S, Cell signaling, 1:100) with 0.5% BSA and 0.3% Triton X-100 in PBS overnight at 4 °C. The retinas were washed in PBS for 30 min at least 8 times at room temperature and incubated with anti-IB4 (I21412, Invitrogen, 1:200) and Alexa Fluor 488 goat anti-rabbit IgG (A11008, Invitrogen, 1:500) as a secondary antibody for eNOS with 1% Triton X-100 in PBS overnight at 4°C in the dark. The retinas were washed in PBS for 30 min at least 8 times at room temperature in the dark, post-fixed in 1% paraformaldehyde in PBS for 5 min and washed with PBS 3 times. The retinas were mounted using Vectashield (Vector Labs) and imaged using fluorescence microscopy.

### Statistical analysis

To analyze significant differences between two groups, two-tailed Student t test or one-way ANOVA was used through GraphPad Prism 9.0. All data were reported as mean ± SEM, and a value of P-value < 0.05 was considered statistically significant.

## RESULTS

### β-arrestins interacts with primary ciliary proteins under shear stress and are required for the formation of primary cilia in HUVECs

β-arrestins 1 and 2 regulate multiple aspects of GPCR signaling, trafficking and endocytosis and are ubiquitously expressed, including in ECs^19^. To identify β-arrestin-interacting partners in ECs under differing hemodynamic states, we exposed human umbilical vein ECs (HUVECs) to either shear stress or a static environment. We then performed proteomics analysis on cell lysates pulled down by β-arrestin1/2 antibody to identify those proteins pulled down under shear stress but not static conditions. Interestingly, we found that β-arrestins bound to several primary ciliary proteins such as IFT81, Kif13A and Kif17 under shear stress, but not static, conditions (**Table 1**). While β-arrestins have been associated with primary cilia in other cell types^14^, their association with cilia in ECs have not been previously demonstrated. Primary cilia are prominent mechanosensors with the ability to physical bend as a result of shear stress. To explore if β-arrestins can co-localize with primary cilia in HUVECs, we transfected a GFP-tagged β-arrestin1 construct to HUVECs and stained cells with the primary cilia marker Ac-α-tubulin. While β-arrestin was detected throughout the cell, we found increased localization of β-arrestin1 to the primary cilia in HUVECs (**Figure 1A**). We further evaluated whether formation of primary cilia in ECs may be affected by disrupted expression of β-arrestins. We found siRNA mediated knockdown of β-arrestin1 and β-arrestin2 led to significant decrease in the percent of ECs with primary cilia (**Figure 1B, C, Supplemental figure 1**), suggesting that β-arrestins play an important role in primary cilia formation or maintenance.

**TABLE 1.**
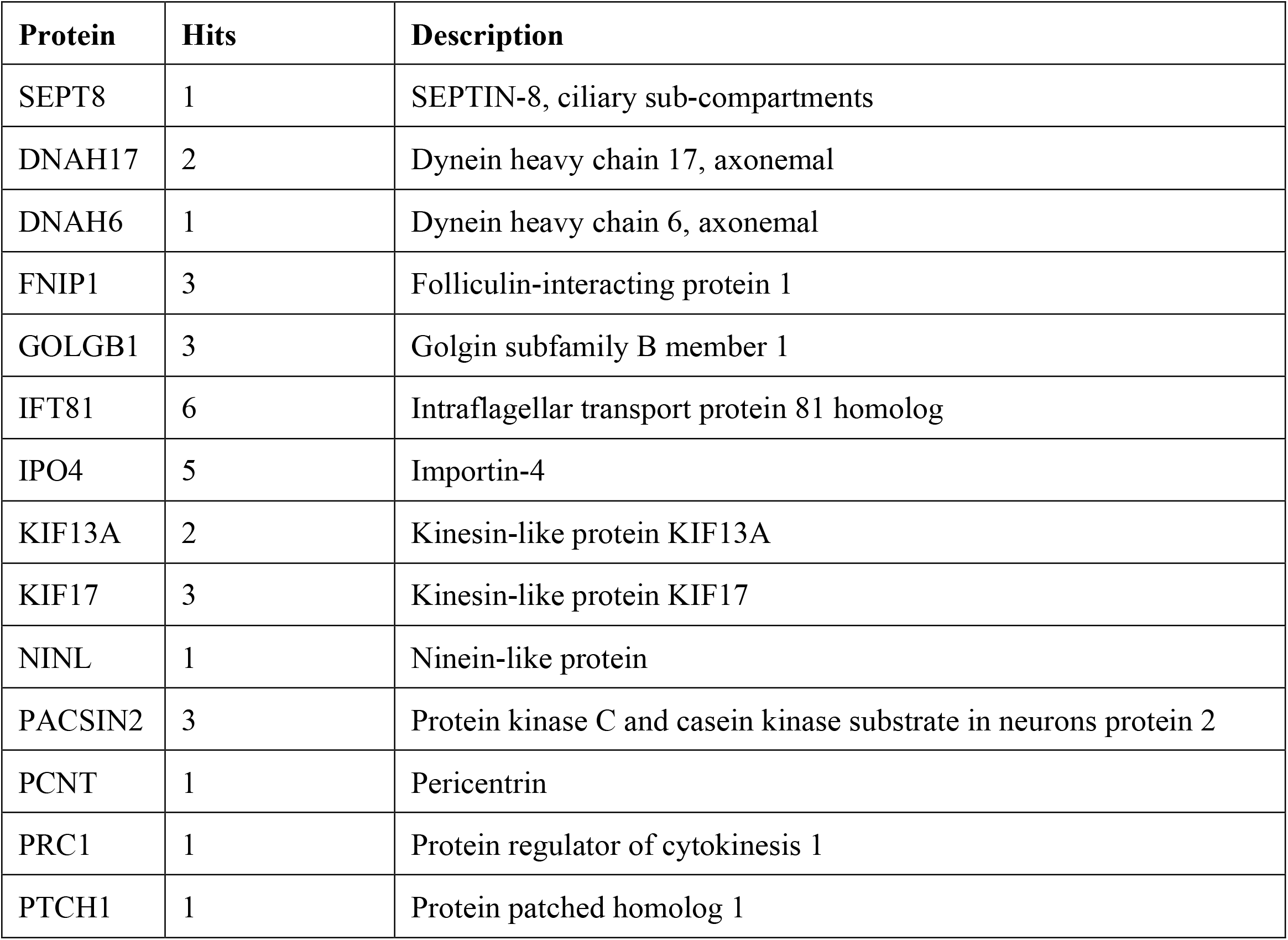
β-arrestin-interacting partners in endothelial cells under shear stress. Human umbilical vein ECs (HUVECs) were exposed to either shear stress or a static environment, and proteomics were performed analysis on cell lysates pulled down by β-arrestin1/2 antibody. Peptides identified under shear stress conditions and not static conditions are listed below.

| Protein | Hits | Description |
| --- | --- | --- |
| SEPT8 | 1 | SEPTIN-8, ciliary sub-compartments |
| DNAH17 | 2 | Dynein heavy chain 17, axonemal |
| DNAH6 | 1 | Dynein heavy chain 6, axonemal |
| FNIP1 | 3 | Folliculin-interacting protein 1 |
| GOLGB1 | 3 | Golgin subfamily B member 1 |
| IFT81 | 6 | Intraflagellar transport protein 81 homolog |
| IPO4 | 5 | Importin-4 |
| KIF13A | 2 | Kinesin-like protein KIF13A |
| KIF17 | 3 | Kinesin-like protein KIF17 |
| NINL | 1 | Ninein-like protein |
| PACSIN2 | 3 | Protein kinase C and casein kinase substrate in neurons protein 2 |
| PCNT | 1 | Pericentrin |
| PRC1 | 1 | Protein regulator of cytokinesis 1 |
| PTCH1 | 1 | Protein patched homolog 1 |

**Figure 1.**
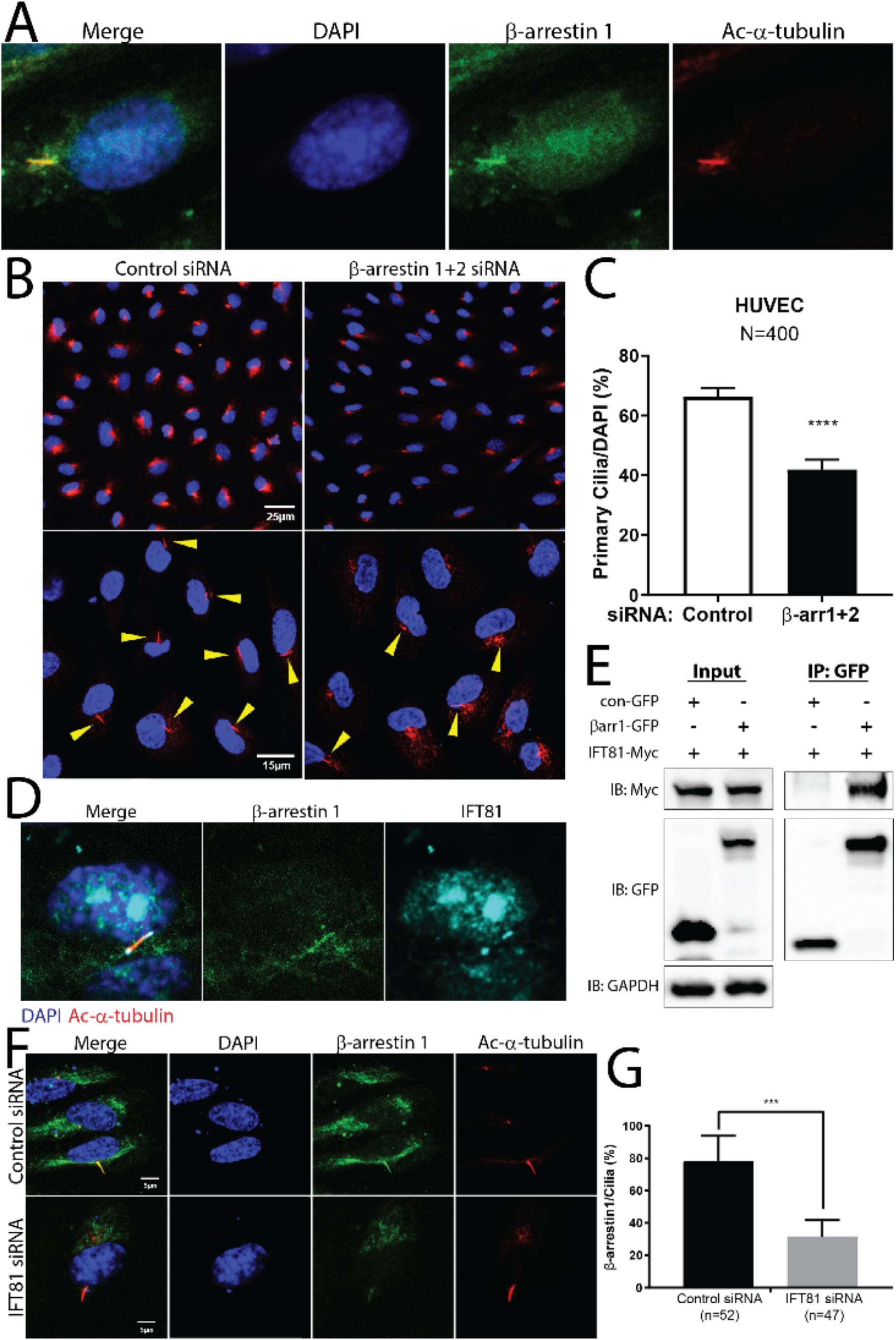
β-arrestin1 interacts with ciliary proteins under shear stress and is required for the formation of primary cilia in HUVECs. **(A)** Immunofluorescence images showing β-arrestin1 (green), primary ciliary marker (red; Ac-α-tubulin) and DAPI (blue) in HUVECs. **(B, C)** Immunofluorescence staining and quantification of primary cilia (red; Ac-α-tubulin) in β-arrestin1/2 siRNA (β-arrs1+2 si) or control siRNA (Con si) treated HUVECs. Yellow arrows indicate primary cilia, and the number of primary cilia was quantified as percentage based on the number of cells (blue; DAPI). N indicates number of cells counted (400) and *\*\*\*\*, p* < 0.0001. **(D)** Immunofluorescence images showing β-arrestin1 (green), IFT81 (cyan), primary ciliary marker (red; Ac-α-tubulin) and DAPI (blue) in HUVECs. **(E)** Immunoprecipitation of GFP-conjugated β-arrestin1 and Myc-conjugated IFT81 in HEK293 cells. Proteins were pulled down with GFP antibody and detected by Myc and GFP antibodies. Control GFP construct (con-GFP) was used as immunoprecipitation control and GAPDH antibody was used as a loading control. **(F, G)** Immunofluorescence staining and quantification of β-arrestin1 (green) in IFT81 siRNA (IFT81 si) or control siRNA (Con si) treated HUVECs. The number of β-arrestin1 localized primary cilia was quantified as percentage based on the number of primary cilia (red; Ac-α-tubulin). n indicates number of primary cilia counted and *\*\*\*, p* < 0.001.

### β-arrestin1 localization to primary cilia is IFT81-dependent

Our proteomics analysis found that IFT81 was the most frequently pulled down protein by β-arrestin1/2 in HUVECs (**Table 1**). IFT81 is a member of the intra-flagellar transport protein family, which is known to be critical in primary cilia formation. IFT proteins are divided into two groups: IFT-A and IFT-B. IFT-A proteins are responsible for retrograde transport with cytoplasmic dynein-2, which involves transport from the ciliary tip back to the cell body. IFT-B proteins are involved in anterograde transport with heterotrimeric kinesin-2, which denotes transport from the base to the ciliary tip. IFT81 belongs to the IFT-B group^26^, and forms a transport complex with IFT74. Previous studies have shown that IFT81 accumulates both on the base and the tip of primary cilia in cells and is involved in elongation of primary cilia^27, 28^.

Given IFT81’s interaction with β-arrestin1/2 and its importance in primary cilia function, we sought to further elucidate the relationship of IFT81 and β-arrestin1/2. We found that IFT81 is co-localized with β-arrestin1 on primary cilia (**Figure 1D**), and directly associates with β-arrestin1 in HUVECs as demonstrated by immunoprecipitation (**Figure 1E**). To assess the role of IFT81 and its interaction with β-arrestin1, we evaluated GFP-tagged β-arrestin1 localization in IFT81-silenced HUVECs. We found that β-arrestin1 localization to the primary cilia was significantly decreased in IFT81-deficient HUVECs (**Figure 1F, G**), suggesting that IFT81 is required for the recruitment of β-arrestin1 to primary cilia. While we expected that the formation of primary cilia would be suppressed in IFT81-deficient HUVECs, we found that IFT81 siRNA knockdown alone did not affect the formation of primary cilia in HUVECs, suggesting that IFT81 may be dispensable for primary cilia formation (data not shown).

### β-arrestins interact with BMPR-II, which is required for the shear stress response

β-arrestins are established downstream signaling partners of GPCRs. With the evidence of β-arrestins being recruited to primary cilia, we next tested whether GPCRs and GPCR cofactors known to be expressed in the vascular endothelium (**Supplemental Table 1**) were localized to the endothelial primary cilia. We found that GFP-tagged S1PR1, CXCR2 and CXCR7 were localized to primary cilia (**Supplemental Figure 1A**), while CXCR1, GPR4, GPR15, PAR1 and RAMP2 did not appear to have preferential ciliary localization (**Supplemental Figure 1B and C**). We next conducted siRNA knockdown of S1PR1, CXCR2 and CXCR7 in ECs to test whether these GPCRs regulated the EC response to shear stress, such as induction of eNOS expression^29^ (**Supplemental Figure 2**). Despite robust knockdown of each GPCR, we did not observe any significant change in eNOS expression or phosphorylation in response to shear stress in ECs (**Supplemental Figure 2**), suggesting that these ciliary GPCRs may not be critical to regulating EC response to shear stress.

Given our findings, we sought to identify additional cell surface receptors that may utilize β-arrestins and be involved in mechanotransduction from endothelial primary cilia. The type II bone morphogenetic protein receptor (BMPR-II) has previously been shown to have a key role in ciliary flow sensing^30, 31^. While β-arrestins have been shown to interact with TGF-β co-receptors^32-34^, whether BMPRs interact with and signal through β-arrestins in the primary cilia has not been determined. We first tested whether β-arrestins are recruited to BMPR-I/BMPR-II complexes (**Figure 2A**). Under basal conditions, β-arrestin2 was found throughout the cytoplasm. After stimulation with BMP-2, β-arrestin2 redistributed to the endosomes, consistent with a ligand-dependent interaction with BMPR-II (**Figure 2A**, right panels).

**Figure 2.**
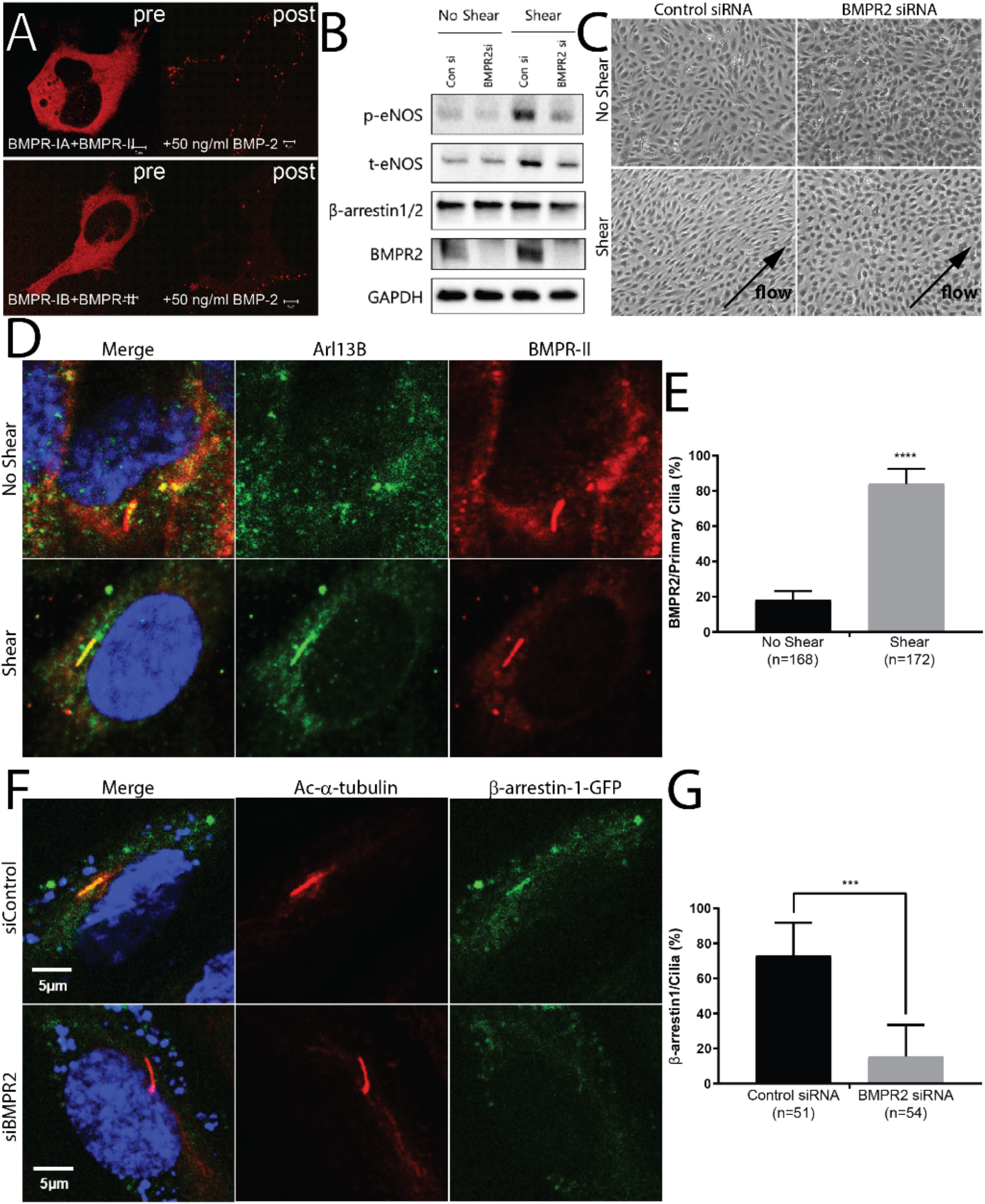
BMPR2 and β-arrestin1 are required for the shear stress response. (**A**) Confocal microscopy demonstrating BMP-2-induced internalization of RFP-labeled β-arrestin2 before (pre) and 30 minutes after stimulation (post) with 50 ng/ml of BMP-2 in HEK293 cells transiently transfected with (top panel) BMPR1A/BMPR2 and (bottom panel) BMPR1B/BMPR2. **(B)** Immunoblot detection of phosphorylated eNOS (p-eNOS), total eNOS (t-eNOS), β-arrestin1/2, BMPR-II and GAPDH levels extracted from BMPR2 siRNA (BMPR2 si) or control siRNA (Con si) treated HUVECs with or without shear stress. **(C)** Cell alignment of BMPR2 siRNA (BMPR2 si) and control siRNA (Con si) treated HUVECs with or without shear stress. **(D, E)** Immunofluorescence staining and quantification of BMPR-II (green) in HUVECs with or without shear stress. The number of BMPR-II localized primary cilia was quantified as percentage based on the number of primary cilia (red; Ac-α-tubulin). n indicates number of primary cilia counted and *\*\*\*\*, p* < 0.0001. **(F, G)** Immunofluorescence staining and quantification of β-arrestin1 (green) in BMPR2 siRNA (si BMPR2) and control siRNA (si Con) treated HUVECs. The number of β-arrestin1 localized primary cilia was quantified as percentage based on the number of primary cilia (red; Ac-α-tubulin). n indicates number of primary cilia counted and *\*\*\*, p* < 0.001.

We further investigated the role of BMPR-II in endothelial shear sensing. We found that eNOS expression under shear stress was decreased in HUVECs subjected to BMPR2 knockdown (**Figure 2B**). Consistent with this finding, physical alignment of cells induced by shear stress was largely absent in BMPR-II-deficient HUVECs (**Figure 2C**). Taken together, these findings are consistent with BMPR-II being critical for induction of eNOS expression and EC alignment in response to shear stress. To investigate whether localization of BMPR-II in primary cilia changed under shear stress, we analyzed the localization of BMPR-II in HUVECs with or without shear stress. Interestingly, we found that under static condition BMPR-II was widely distributed throughout the cell, but subjecting ECs to shear stress led to significant increase in BMPR-II localization to the primary cilia (**Figure 2D, E**). We next assessed whether BMPR-II may be involved in recruitment of β-arrestin1 to the primary cilia. We transfected GFP-tagged β-arrestin1 constructs to BMPR2-knockdown HUVECs and stained for primary cilia. We found that the localization of β-arrestin1 to primary cilia was markedly decreased in BMPR2-silenced HUVECs (**Figure 2F, G**). Our results demonstrate the importance of BMPR-II in regulation of shear stress response in ECs and its role in regulating β-arrestin translocation to the primary cilia.

### β-arrestins interact with and regulate signaling by BMPR-II

We next tested whether β-arrestins could directly interact with BMPR-II and regulate its signaling. Co-immunoprecipitation of overexpressed BMPR-II and β-arrestins in HEK293 cells demonstrated a constitutive interaction between BMPR-II and β-arrestin1 and 2 (**Figure 3A**); this interaction was preserved in a BMPR2 truncation mutant (S532X) that removed the long C-terminal tail after the S/T kinase domain (**Figure 3A**). In the context of confocal microscopy that demonstrated β-arrestin redistribution after ligand stimulation (**Figure 2A**), this is consistent with β-arrestins interacting with BMPR-II in both a constitutive and ligand-induced manner. We next examined the effects of loss of β-arrestin1 and 2 on BMPR-II signaling as assessed by Smad1/5/8 (Smad) phosphorylation. HEK293 cells were transfected with β-arrestin1 or β-arrestin2 siRNA and stimulated with BMP-2. We observed that β-arrestin1 knockdown significantly decreased BMP-induced Smad phosphorylation while β-arrestin2 knockdown had no effect (**Figure 3B**), except for a modest increase at 15 minutes (**Figure 3C**). We then tested whether β-arrestin1 or 2 siRNA knockdown had any effect of transcription of a luciferase reporter with BMP-responsive elements from the *Id1* promoter, providing a downstream readout of BMPR-II signaling^21^. Although β-arrestin1 and β-arrestin2 had different effects on Smad phosphorylation, knockdown of either significantly decreased the activity of this reporter transfected in HEK293 cells stimulated with BMP-2 (**Figure 3D**). Taken together, these data demonstrate that β-arrestins regulate Smad phosphorylation by BMPR2 and its transcriptional activity.

**Figure 3.**
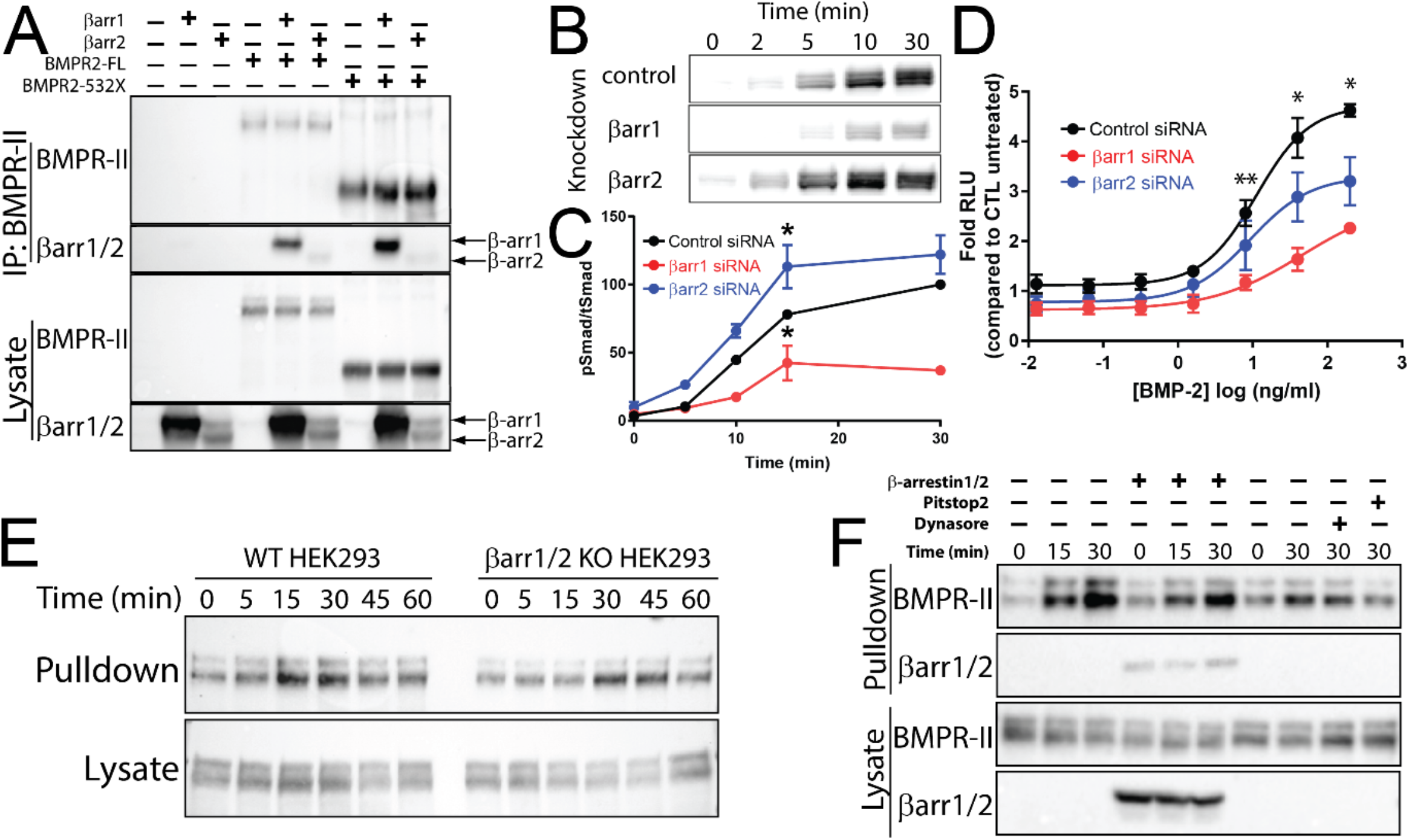
β-arrestins interact with and regulate signaling by BMPR-II. (**A**) Interaction of β-arrestins with BMPR-II as assessed by co-immunoprecipitation of overexpressed FLAG-tagged β-arrestin1/2 with myc-tagged full length BMPR2 and its 532X truncation. (**B, C**) β-arrestin1 significantly reduces BMP-induced Smad1/5/8 phosphorylation by BMPR-II after stimulation with BMP-2 (50 ng/ml) with siRNA knockdown of β-arrestin1 or β-arrestin2. pSmad/tSmad levels were normalized to ctl siRNA signal at thirty minutes. (*, p < 0.05 compared to other siRNA treated samples for that time point by two-way ANOVA with Tukey’s multiple comparison test) (**D**) Both β-arrestin1 and 2 knockdown significantly decreases activity of a BMP reporter (Bre-luc) transfected in HEK293 cells stimulated with BMP-2 (10 ng/ml). (*, p < 0.05 all groups significantly different; **, p < 0.05 β-arrestin1- and ctl-siRNA significantly different by two-way ANOVA with Tukey’s multiple comparison test). (**E**) No difference in time course of BMP-2-induced endocytosis between WT and β-arrestin1/2 KO HEK293 cells. The endocytosed pool of BMPR-II was labeled with biotin (see *Methods*) prior to pulldown with avidin beads. (**F**) No effect of β-arrestin1/2 overexpression on receptor internalization in β-arrestin1/2 KO cells. Endocytosis at 30 minutes was significantly decreased by the clathrin inhibitor Pitstop2 but not by the dynamin inhibitor dynasore. All experiments performed at least 3 times; shown are mean ± SEM. Blots and images shown are representative from at least 3 independent experiments.

For many GPCRs, β-arrestins act as adaptor proteins for receptor endocytosis. We tested for such a role at BMPR2 by assessing receptor endocytosis in combination with either β-arrestin1/2 knockout (KO) by CRISPR/Cas9^35^ or overexpression of β-arrestin1/2. Using a receptor biotinylation assay to selectively pulldown endocytosed receptor^36^, we did not observe any change in receptor internalization in a β-arrestin1/2 KO HEK293 cell line compared to the parental cell line (**Figure 3E**). Restoration of β-arrestin1/2 expression in the KO cells did not impact internalization (**Figure 3F**), consistent with β-arrestins not playing a role in BMP-induced receptor internalization. Receptor internalization did decrease significantly with clathrin inhibition, but was not affected by dynamin inhibition (**Figure 3F**). These findings are consistent with ligand-induced BMPR-II internalization requiring clathrin but neither β-arrestins nor dynamin.

### β-arrestin is required for the EC response to shear stress and eNOS expression

To investigate whether β-arrestin is required for EC signal transduction in response to shear stress, we tested for changes in HUVEC orientation and eNOS expression with or without shear stress in β-arrestin-deficient HUVECs. HUVEC cell alignment in response to shear stress was maintained in control siRNA treated cells but were lost in both β-arrestin1-or 1+2 siRNA-treated cells (**Figure 4A**). Consistent with this, we found that induction of eNOS expression in response to shear stress was lost in HUVECs subjected to β-arrestin knockdown (**Figure 4B, C**). These results were consistent with β-arrestins being required for the EC response to shear stress.

**Figure 4.**
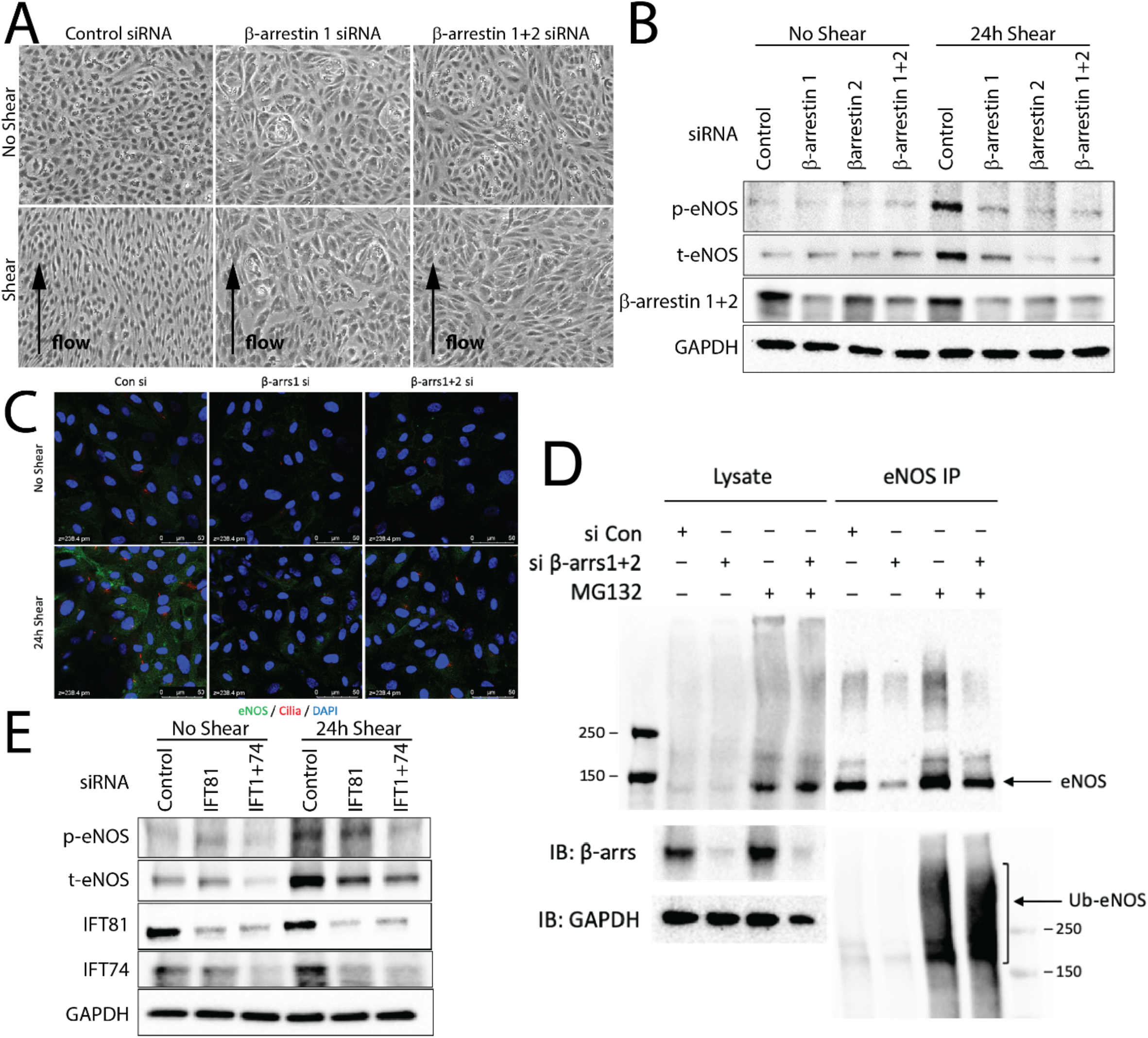
β-arrestin1 is required for the endothelial cell shear stress response and eNOS expression. **(A)**Cell alignment of β-arrestin1/2 siRNA or control siRNA treated HUVECs with or without shear stress. **(B)**Immunoblot detection of phosphorylated eNOS (p-eNOS), total eNOS (t-eNOS), β-arrestin1/2 and GAPDH levels extracted from β-arrestin1/2 siRNA or control siRNA treated HUVECs with or without shear stress. **(C)** Immunofluorescence images showing eNOS (green) and DAPI (blue) in β-arrestin1/2 siRNA or control siRNA treated HUVECs with or without shear stress. **(D)** Immunoprecipitation of ubiquitin-eNOS in β-arrestin1/2 siRNA treated HUVECs. Proteins were pulled down with eNOS antibody and detected by ubiquitin-eNOS and total eNOS antibodies. Input lysates were detected by β-arrestin1 and GAPDH antibodies. MG132 is a proteosomal inhibitor. **(E)** Immunoblot detection of phosphorylated eNOS (p-eNOS), total eNOS (t-eNOS), IFT81, IFT74 and GAPDH levels extracted from IFT81 and/or IFT74 siRNA or control siRNA treated HUVECs with or without shear stress.

We next sought to determine the mechanism of disrupted shear response in β-arrestin deficient ECs. First, we found that while eNOS protein levels were not upregulated in response to shear stress in β-arrestin knockdown ECs, the increase in eNOS mRNA transcript by shear stress was not significantly affected by β-arrestin knockdown (**Supplemental Figure 3)**. Moreover, transcription factors Krüppel-like Factor 2 (KLF2) and Krüppel-like Factor 4 (KLF4), which are well known to be transcriptionally induced by shear stress and regulate eNOS expression,^37^ were also not affected by β-arrestin knockdown (**Supplemental Figure 3**). These findings suggest that shear stress induced eNOS transcriptional upregulation is not affected by β-arrestin knockdown. To further explore the mechanism of β-arrestins in modulating eNOS protein expression, we investigated ubiquitination of eNOS. eNOS is regulated by Hsp90-based chaperone mediated ubiquitin-proteasome system^38, 39^. It is reported that MG132, which is a proteosomal inhibitor, increases eNOS expression and NO generation in HUVEC^40^. Interestingly, we found that ubiquitination of eNOS was increased in response to β-arrestin knockdown (**Figure 4D**), which demonstrated that β-arrestins contributes to the shear response in HUVECs by inhibiting degradation of eNOS. We further evaluated whether ciliary proteins, namely the IFT proteins, can directly modulate eNOS induction in response to shear stress. We found that shear stress induced eNOS expression was significantly decreased in response to IFT81 knockdown, which was further decreased via IFT81 and IFT74 combined knockdown in HUVECs (**Figure 4E**). These findings are consistent with ciliary structural proteins being required for β-arrestin-mediated regulation of eNOS expression in HUVECs.

### Retinal vascular development is impaired in Arrb1/Arrb2 ECKO mice

Primary cilia play a key role in vascular development, with expression in mouse retinal endothelia where they promote vessel stabilization through increased sensitivity of the BMP-Alk1 pathway^17^. We generated endothelial-specific, conditional β-arrestin1 and β-arrestin2 double knockout mice (*Arrb1*/*Arrb2* ECKO mice) to determine whether β-arrestins could affect primary cilia and vascular development. Mouse pups were injected with tamoxifen on postnatal day (P) 1 through P3 to induce endothelial *Arrb1/Arrb2* deletion and were analyzed at P7 to assess for changes in retinal vascular development. We found that vascular plexus of *Arrb1/Arrb2* ECKO retinas was markedly reduced compared to that of control retinas (**Figure 5A**) with shorter radial expansion length compared to that of control retinas (**Figure 5B**), and a significant reduction in the number of branchpoints was observed in *Arrb1/Arrb2* ECKO retinas compared to controls (**Figure 5C**). These findings suggest that endothelial β-arrestin1 and β-arrestin2 play an important role in vascular development. Notably, mouse retina vessels have been reported to have different numbers of primary cilia depending on the specific vessel bed, with increased numbers in the veins and plexus, and fewer in the arteries^31^. Therefore, we tested whether there was a difference in the number of primary cilia between veins and arteries in mice retina and whether this differed in *Arrb1/Arrb2* ECKO compared to wild-type retina. We quantified the number of primary cilia and normalized them to the area of vessels. We found no difference in the number of primary cilia between different vessels, nor any significant difference between *Arrb1/Arrb2* ECKO and wild-type retinas (**Supplemental Figure 4**). To determine whether β-arrestin also regulates eNOS expression *in vivo* as we observed *in vitro*, we stained *Arrb1/Arrb2* ECKO retinas for eNOS. Consistent with our *in vitro* findings, *Arrb1/Arrb2* ECKO retinas displayed significant decrease in eNOS expression compared to control retinas, consistent with β-arrestins being critical for eNOS expression in the vasculature *in vivo* (**Figure 5D**).

**Figure 5.**
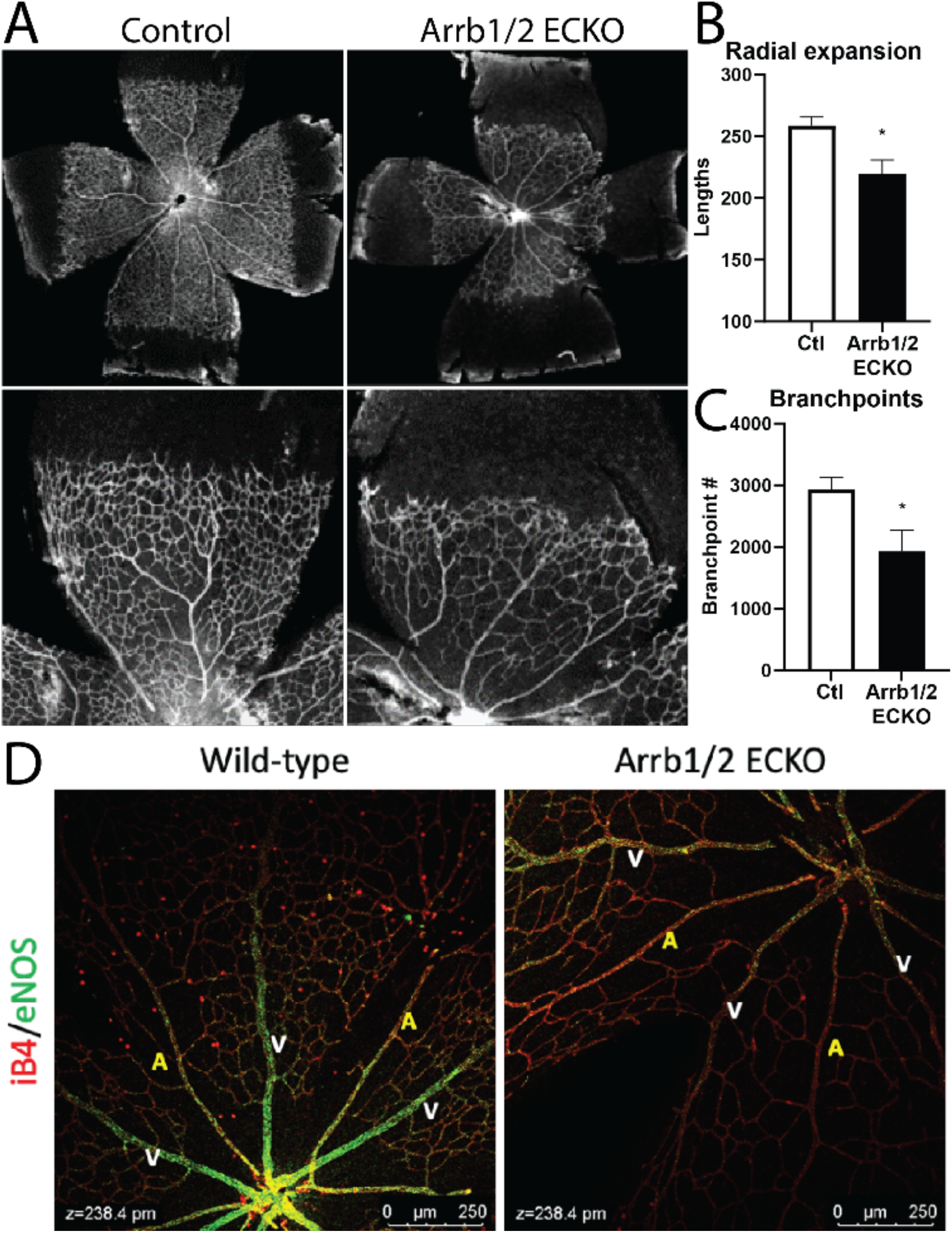
Endothelial-specific knockout of β-arrestins results in a retinal vascular development defect. **(A)** IB4 stained p7 retinal flat-mounts of *Arrb1/Arrb2* ECKO and wild-type (Control) mice. **(B)** Quantification of vascular radial expansion in *Arrb1/Arrb2* ECKO and wild-type (Ctl) mice. N = 7 wild-type and 4 Arrb1/2 ECKO. *\*, p* < 0.05. **(C)** The number of branchpoints in *Arrb1/Arrb2* ECKO and wild-type (Ctl) mice. N = 7 wild-type and 3 Arrb1/2 ECKO. *\*, p* < 0.05. **(D)** eNOS (green) and IB4 (red) stained p7 retinal flat-mounts of Arrb1/2 ECKO and wild-type mice. A indicates arteries and V indicates veins.

## DISCUSSION

The primary cilium is a sensory organelle that receives both mechanical and chemical signals from other cells and the environment. The integration of these different stimuli through primary cilia is central to many aspects of endothelial function^8, 41^. Primary cilia are present in almost all types of cells and essential signal transducers that dictate various cellular responses. For example, multiple components of the sonic hedgehog signaling cascade and many growth factors localize to the primary cilium, and loss of the cilium blocks ligand-induced signaling, leading to uncontrollable cell growth or disruptions in homeostasis^42^. Primary cilia are largely absent in high laminar flow, athero-resistant regions of vessels but found in regions of disturbed flow^43^ or areas of high shear stress^44^. Mechanosensing responses of primary cilia in ECs play important roles in many vascular functions such as in the pulmonary circulation^41^. Here we identified a novel signaling mechanism in which β-arrestins regulate ciliary BMPR-II signaling that contributes to the endothelial response to shear stress.

We found that β-arrestins, already known to regulate signal transduction, are required for formation and mechanosensing responses of primary cilia in ECs. β-arrestins associated with IFT81 to localize in the primary cilia and this complex likely plays role in formation of the primary cilia, as β-arrestin knockdown decreased the number of primary cilia. We also found that the eNOS-mediated shear response via primary cilia in ECs was regulated by β-arrestins and required BMPR2. We also demonstrated that β-arrestins were required for BMPR-II signaling through phosphorylation of Smad transcription factors, consistent with their role in regulating the endothelial mechanosensing response. While β-arrestins have previously been shown to play a role in regulating signaling by TGF-β coreceptors ^32, 33^, our study identified a role for β-arrestins in promoting signaling by a type II TGF-β receptor, BMPR-II. Notably, while β-arrestins regulate multiple aspects of GPCR signaling and endocytosis, we only found that β-arrestins were required for Smad phosphorylation and the downstream transcriptional response but not required for receptor endocytosis. While we did not elucidate the full mechanistic details that underlie β-arrestin regulation of BMPR-II, this suggests that the interaction of β-arrestins with BMPR-II relies on different mechanisms than β-arrestin regulation of GPCRs.

With the well-described role of β-arrestins in regulating GPCR signaling^19^, we attempted to find GPCRs which regulate endothelial shear response via interacting with β-arrestins in primary cilia based on the fact that β-arrestin is a multifunctional regulator for most of GPCRs. We had 8 potential GPCR candidates that are known to affect vascular development and functions, however, the candidates we tested did not appear to promote eNOS-mediated endothelial shear response through β-arrestins. It is likely that other GPCRs that are localized in primary cilium interact with β-arrestins to regulate endothelial mechanosensing^45-47^. Based on the localization of other GPCRs, including S1PR1, CXCR2 and CXCR7, β-arrestins may play a role in integrating signals from a number of receptors in the primary cilium that regulate the endothelial response to shear stress.

Endothelial mechanosensing plays an important role not only in the response to shear stress in the adult, but also in vascular development^12^. To address whether β-arrestins played a role in vascular development, we generated mice with inducible, endothelial specific knockout of both β-arrestins (*Arrb1/Arrb2* ECKO mice). We generated these mice because β-arrestin1 homozygous knockout mice have essentially normal basal cardiovascular physiology including heart rate, blood pressure and left ventricular ejection fraction but display an abnormal response to β-adrenergic stimulation^48^. Similarly, β-arrestin2 knockout mice are phenotypically normal and produce viable progeny but display abnormal responses to physiologic stresses^49^. In contrast, while mice deficient of either β-arrestin1 or β-arrestin2 develop normally and are fertile, β-arrestin1 and β-arrestin2 double knockout mice die within few hours after birth due to pulmonary hypoplasia^50^. This is consistent with β-arrestins compensating for one another in the setting of knockout of the other isoform. *Arrb1/Arrb2* ECKO mice allowed us to determine whether changes in the EC shear stress response mediated by β-arrestins result in a vascular phenotype. We found that *Arrb1/Arrb2* ECKO mice displayed abnormal retinal vascular development, with decreased radial expansion and a loss of branchpoints. Consistent with this, we found that *Arrb1/Arrb2* ECKO retinas displayed significantly less eNOS localization at arteries and veins than WT retinas. These findings are consistent with β-arrestins regulating the endothelial shear response through BMPR2, but may also reflect a role for β-arrestins in regulating GPCRs that are important in vascular development.

In conclusion, we report a novel function of endothelial β-arrestins in primary ciliary mechanosensing via BMPR-II signaling. Our finding contribute to an enhanced understanding of the regulation of ciliary signaling and how flow-induced shear stress regulates receptor signaling and vascular development. Future studies will be required to address the detailed biochemical mechanisms that underlie β-arrestin regulation of BMPR-II signaling and how signaling through different receptors in the primary cilia is integrated through β-arrestins and other signaling proteins.

## Nonstandard Abbreviations and Acronyms

*Arrb1/Arrb2*: ECKO Endothelial specific, conditional β-arrestin1 and β-arrestin2 double knock-out
BMPR-II: Type II bone morphogenetic protein receptor (protein)
BMPR2: Type II bone morphogenetic protein receptor (gene)
eNOS: Endothelial nitric oxide synthase
GPCR: G protein-coupled receptor
HUVEC: Human umbilical vein endothelial cell
IFT81: Intraflagellar transport protein 81

## ACKNOWLEDGEMENTS

We thank Jeffrey Kovacs for valuable discussions and help with BMPR2 biochemical studies. We thank Gerard Blobe and Robert Lefkowitz for valuable discussions. We thank Asuka Inoue for the β-arrestin1/2 KO HEK293 cells, Peter ten Dijke for the *bre-luc* reporter construct and Nicholas Morrell for the *BMPR2* S532X mutant construct.

