## Supplemental data for "Endothelial β-arrestins Regulate Mechanotransduction by the Type II Bone Morphogenetic Protein Receptor in Primary Cilia"

**Supplemental Table 1. G protein-coupled receptors and related proteins assessed for localization in primary cilia in endothelial cells.**

| Receptor | Function / Phenotype | References |
|---|---|---|
| **S1PR1**<br>sphingosine-1-phosphate receptors, EDG1 | • S1P associates with the S1PR1 and regulates angiogenesis and oncogenesis<br>• Genetic disruption of a single S1pr1 gene causes vascular defects | Yu H. et al., 2015, Genes Cells, 20(8):647-58[1]<br>Karen M. et al., 2013, JBC, 288(4): 2143-2156[2] |
| **CXCR2**<br>C-X-C motif chemokine receptor2 | • Receptors for angiogenic CXC chemokines<br>• CXCR2 silencing modulates CXCL8-dependent endothelial capillary-like structure formation | Seema S. et al., 2011, Microvasc Res, 82(3): 318-25[3] |
| **CXCR7**<br>Atypical chemokine receptor 3, ACKR3, C-X-C chemokine receptor type 7 | • Loss of endothelial CXCR7 reduced vascular density<br>• CXCR7 interacts with β-arrestins and are responsible for G-protein-independent signals through ERK1/2 phosphorylation | Circulation, 2017, 135:1253-1264[4]<br>PLoS One, 2012, 7(3):e34192[5] |
| **CXCR1**<br>C-X-C motif chemokine receptor1 | • Receptors for angiogenic CXC chemokines<br>• Silencing of CXCR1 inhibited capillary-like structure formation by reducing stress fibers | Seema S. et al., 2011, Microvasc Res, 82(3): 318-25[3] |
| **GPR4**<br>G-protein-coupled receptor4 | • GPR4 functions as a pH sensor and GPR4-defected mice have vascular abnormalities<br>• Mice lacking GPR4,proton sensing receptor, reduces pathological angiogenesis | Li V. et al., 2007, Mol Cell Biol.[6]<br>Wyder L. et al., 2011, Angiogenesis[7] |
| **GPR15**<br>G-protein-coupled receptor15 | • GPR15 mediates angiogenesis and cytoprotective function of thrombomodulin | Pan B., et al., 2017, Sci Rep., 7(1):692[8] |
| **PAR1**<br>Proteinase-activated receptor1 | • PAR1 is expressed in vascular cells and is involved in atherosclerosis | Nikos E., et al., 2007, Semin Thromb Hemost[9] |
| **RAMP2**<br>Receptor activity modifying protein2 | • GPCR modulator protein RAMP2 is essential for angiogenesis and vascular integrity<br>• In RAMP2 -/- mice, Vascular abnormalities and reduced responses to angiogenic stimuli | Yuka I., et al., 2008, J Clin Invest 118[10] |

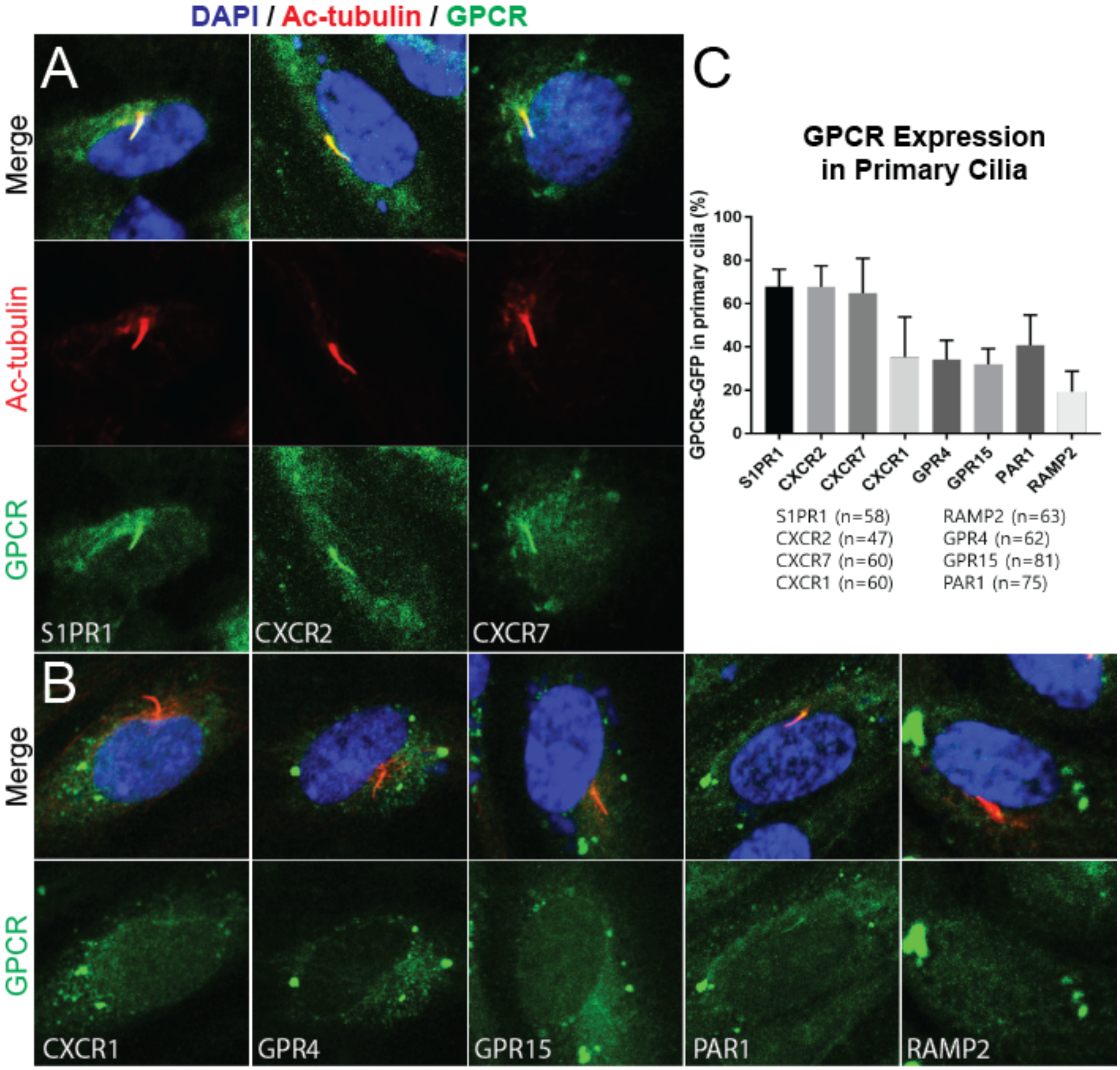


**Supplemental Figure 1. GPCR expression in endothelial cell primary cilia.** (**A**) S1PR1, CXCR2 and CXCR7 (green) are expressed in the primary cilia (Ac-tubulin, red) in the merged image (DAPI, blue). (**B**) No colocalization is observed for other candidate GPCRs and other transmembrane proteins that were tested: CXCR1, GPR4, GPR15, PAR1 and RAMP2. (**C**) Quantification of GPCR GFP signal in primary cilia demonstrates that S1PR1, CXCR2 and CXCR7 are expressed in the primary cilia, while the others do not.

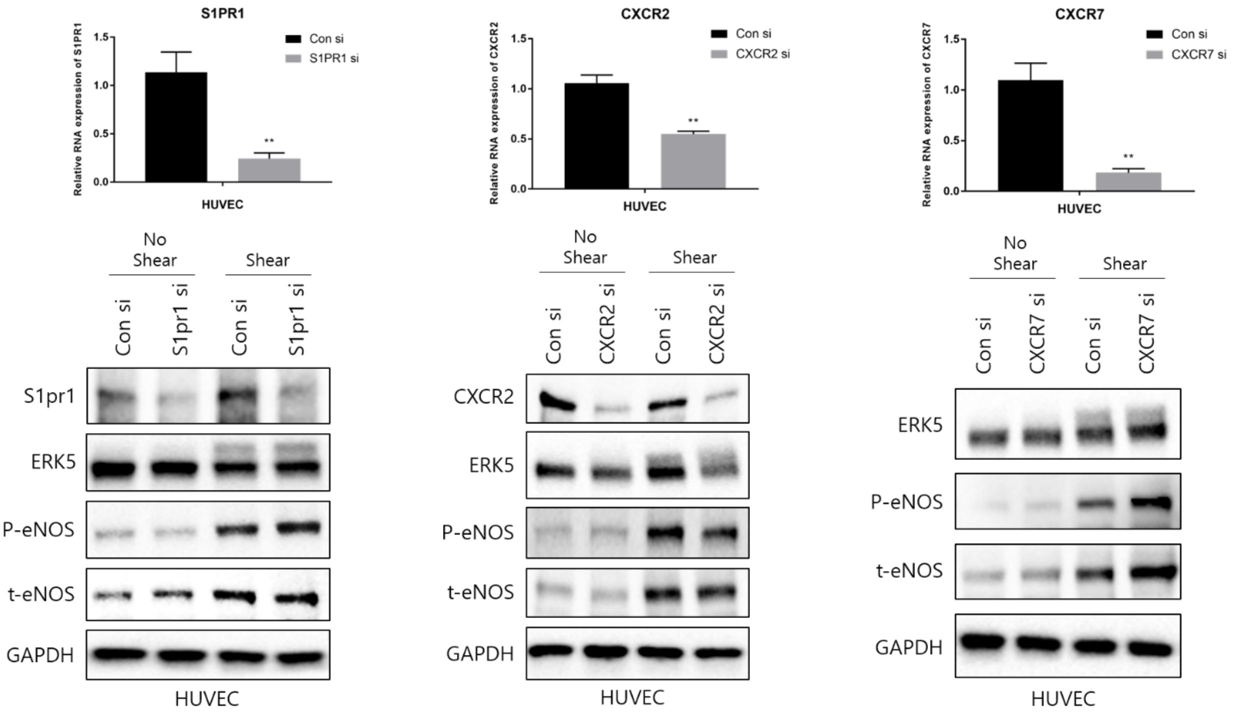


**Supplemental Figure 2. Ciliary GPCR knockdown has no significant effect on eNOS phosphorylation in response to shear.** (**Top panels**) Efficiency of GPCR knockdown. (**Bottom panels**) Knockdown of S1PR1, CXCR3 and CXCR7 had no effect on eNOS phosphorylation in response to shear stress.

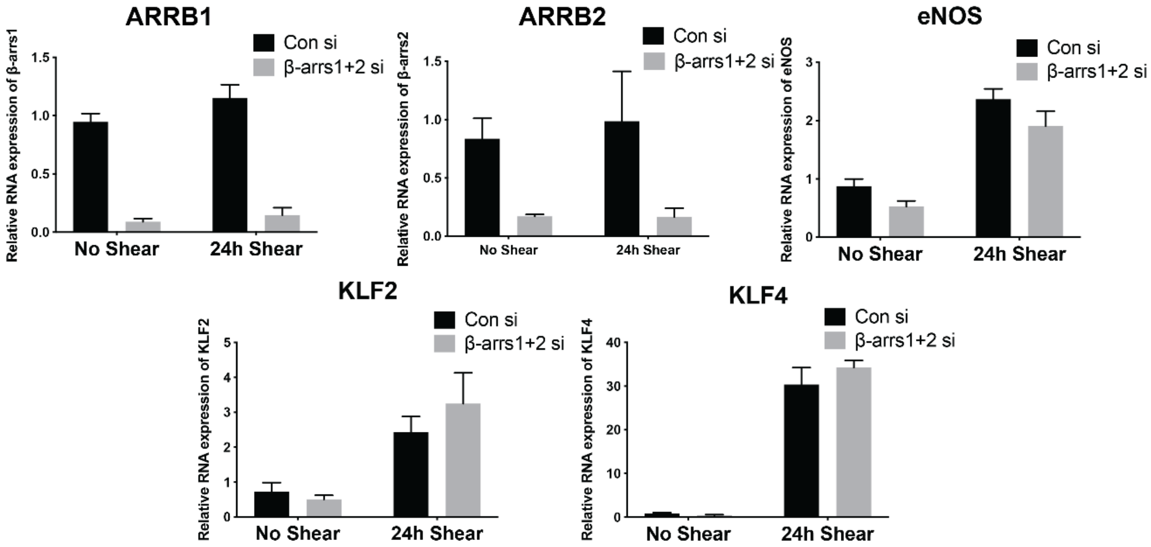


**Supplemental Figure 3. qPCR of targets in response to shear stress in HUVECs.**

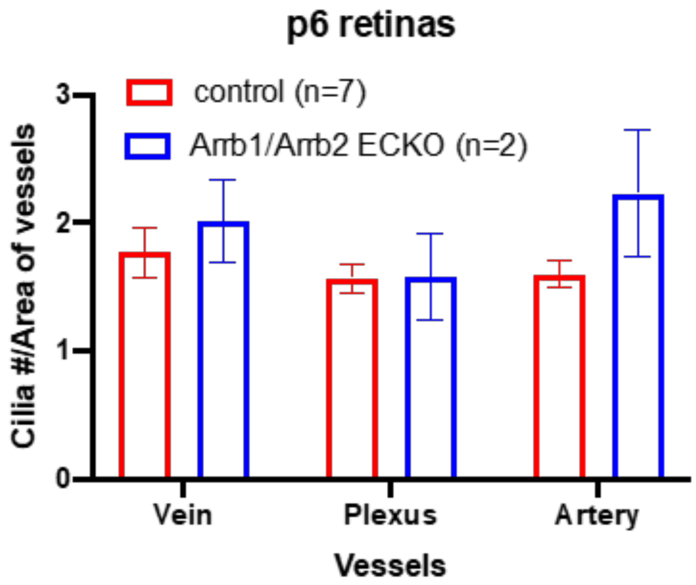


**Supplemental Figure 4.** The number of primary cilia did not differ significantly between vein, plexus, and artery or between control and Arrb1/Arrb2 ECKO retinas.
